# α-Catenin Dependent E-cadherin Adhesion Dynamics as Revealed by an Accelerated Force Ramp

**DOI:** 10.1101/2023.07.28.550975

**Authors:** Joshua Bush, Jolene I. Cabe, Daniel Conway, Venkat Maruthamuthu

## Abstract

Tissue remodeling and shape changes often rely on force-induced cell rearrangements occurring via cell-cell contact dynamics. Epithelial cell-cell contact shape changes are particularly dependent upon E-cadherin adhesion dynamics which are directly influenced by cell-generated and external forces. While both the mobility of E-cadherin adhesions and their adhesion strength have been reported before, it is not clear how these two aspects of E-cadherin adhesion dynamics are related. Here, using magnetic pulling cytometry, we applied an accelerated force ramp on the E-cadherin adhesion between an E-cadherin-coated magnetic microbead and an epithelial cell to ascertain this relationship. Our approach enables the determination of the adhesion strength and force-dependent mobility of individual adhesions, which revealed a direct correlation between these key characteristics. Since α-catenin has previously been reported to play a role in both E-cadherin mobility and adhesion strength when studied independently, we also probed epithelial cells in which α-catenin has been knocked out. We found that, in the absence of α-catenin, E-cadherin adhesions not only had lower adhesion strength, as expected, but were also more mobile. We observed that α-catenin was required for the recovery of strained cell-cell contacts and propose that the adhesion strength and force-dependent mobility of E-cadherin adhesions act in tandem to regulate cell-cell contact homeostasis. Our approach introduces a method which relates the force-dependent adhesion mobility to adhesion strength and highlights the morphological role played by α-catenin in E-cadherin adhesion dynamics.

## Introduction

Cells possess a mechanical nature which is inherently viscoelastic – however they display unique nonlinear mechanical properties which resemble that of glassy polymers and semiflexible networks rather than classical viscoelastic materials which can be described as simple spring-dashpot systems [1-7]. There are a range of measurement techniques which can yield high-resolution localized measurements via contact methods such as nanoindentation [8] or active/passive microrheology [2, 9-11]. Alternatively, non-contact methods enable unperturbed spatial mapping of cell mechanical properties via ultrasound elastography or optical elastography [12-16]. However, contact methods permit the study of force-responsive events such as stiffening or fluidization [17-19], as well as the isolation of passive cell mechanics from the forces generated within the cell itself [20, 21]. Magnetic tweezers can be used to twist a cell-bound magnetic bead for adhesion specific rheology [22] Alternatively, the compliance of the bead in response to a pulling force can be used to independently extract the rheological properties [23, 24], or by applying a great enough force to remove the bead from the cell, the adhesion strength [25]. In this study, we introduce a simple new protocol which enables us to simultaneously probe both the force-dependent mobility and adhesion strength of cell-cell adhesions by using a magnetic pulling cytometer to progressively pull on a cell-bound bead. Specifically, we apply an accelerated force ramp, inspired by the progressive pulsatile force-generation of the actomyosin network [26-31], to correlate the active response mechanics of E-cadherin adhesion with morphological observations of cell-cell contacts.

During morphogenesis and adult tissue remodeling, tissue level shape changes are ubiquitous. Such changes often involve smaller scale cell shape changes that occur in the presence of cell-cell contacts. For instance, apical constriction of cells underlies tissue invagination during gastrulation [32]. Cell-cell contacts in epithelia are mediated by multiple adhesion systems, including tight junctions, desmosomal junctions and adherens junctions. In particular, E-cadherin is a major constituent of adherens junctions which help organize other junctions and actively respond to mechanical forces. E-cadherin adhesions are defined as micron-scale collectives of *trans* E-cadherin bonds formed by the binding of the extra-cellular regions of E-cadherin from neighboring cells, and couple to the cytoskeleton [33] via a host of intermediaries recruited at their cytoplasmic ends. They thereby bind neighboring cells at the cell-cell contact and are subject to both internally cell-generated and externally applied forces. Individual micron-scale E-cadherin adhesions have been shown to displace while still maintaining adhesion [34]. Such displacement can thus be expected due to cell-generated forces due to actomyosin contractility as well as external forces exerted by neighboring cells, that also ultimately arise from actomyosin contractility. How E-cadherin adhesions resist and otherwise displace in response to forces, and at what force levels they rupture, determines how cell-cell contacts evolve with time and thereby effect tissue-level shape changes.

E-cadherin adhesions primarily experience tension [35-38] via their coupling to the actin cytoskeleton through the catenins [39-41]. Specifically, E-cadherin binds to β-catenin which in turn binds to α-catenin [42]. α-catenin couples both directly to F-actin and via a host of potential intermediaries [43, 44], including vinculin [45], ZO-1 [46], afadin [47], α-actinin [48], Fmn1 [49] and EPLIN [50]. As a mechanotransducer, α-catenin unfolds in response to force and recruits vinculin [51, 52] and displays independent mechanotransducing behaviors such as opposing slip and catch bond behaviors [52, 53] as well as force-induced dimerization [41]. Specifically, α-catenin has been shown to be required for the high adhesion strength of single E-cadherin-E-cadherin linkages [54] as well as the cell-cell interface between cells expressing E-cadherin [55, 56], with the α-catenin-actin link essential for strengthening of the cell-cell contact. The mobility of E-cadherin adhesions within the cell-cell contact has also been shown to vary dependent upon α-catenin [57]. However, whether there is an α-catenin-dependent relation between E-cadherin adhesion strength and force-dependent mobility remains unclear.

Adhesion strength measurements of E-cadherin have typically been performed either at the single molecule level [54, 58] or whole cell level [55, 59]. There is a relative lack of approaches that determine E-cadherin adhesion strength at the micron-scale adhesion level. Measurements at the adhesion-scale contain information on functional adhesive units that are not present in molecular measurements. For instance, localized lateral contractions between E-cadherin adhesions within the same cell have been shown to play a role in E-cadherin-coated substrate rigidity sensing [30]. Adhesion-scale measurements also avoid confounding factors such as specific overall geometry and the spatiotemporal experience of force which differs from that of native cell-cell contacts in some cell-scale measurements [59]. While the response of E-cadherin-coated beads adherent to cells and subject to oscillatory forces [60, 61], such as stiffening via mechanotransduction, has been reported [62], adhesion between cells often experience sustained pulling forces. Furthermore, how adhesion mobility is altered under force before rupture is less clear. We therefore sought the force response of an E-cadherin adhesion, along with the adhesion strength of the same adhesion, by exerting an accelerated force ramp on an E-cadherin coated microbead bound to an epithelial cell. We found that E-cadherin adhesion strength is correlated with the adhesion’s mobility in response to force, as quantified by the effective drag coefficient before rupture. We also found that, in the absence of α-catenin both the effective drag coefficient and adhesion strength decreased, along with the diminished potential for cell-cell contacts to recover from a fibrous state when strained.

## Materials and Methods

### Cell culture

Madin-Darby Canine Kidney (MDCK II) cells were grown in DMEM (Dulbecco’s modified Eagle’s medium, Corning Inc., Corning, NY) mixed with L-Glutamine, sodium pyruvate, 1% Penicillin/Streptomycin and 10% Fetal Bovine Serum (FBS) (Corning Inc., Corning, NY), at 37 °C under 5% CO_2_. To generate MDCK α-catenin knockout (KO) cells, CRISPR/Cas9 was used with the gRNA sequence TCTGGCAGTTGAAAGACTGT as employed previously [63]. Cells were transiently transfected with the Sigma All-in-One U6-gRNA/CMV-Cas9-tGFP Vector (with the gRNA), followed by clonal expansion. Cells were plated on No.1.5 glass coverslips coated with Collagen-1 (Dow Corning). The coverslip was prepared by a 5 min exposure to deep UV light (Novascan Technologies, Boone, IA) and a subsequent 30 min incubation with 0.1 mg/mL Collagen-1 in a Phosphate Buffered Saline (PBS) solution. The prepared coverslips were stored at 4°C in PBS for up to 24 hours. Approximately 10^4^ cells were plated onto the Col-1 coated coverslip and incubated in cell culture media at 37°C overnight.

### Imaging

A DMi8 epifluorescence microscope (Leica Microsystems, Buffalo Grove, IL) equipped with a Clara cooled CCD camera (Andor Technology, Belfast, Ulster, UK) was used to obtain phase and fluorescence images.

### Magnetic Bead Coating

Carboxylate functionalized 2.8 μm superparamagnetic beads (M-280 Dynabeads, Thermo Fisher Scientific, Waltham, MA) were coated with protein A (Prospec, East Brunswick, NJ) and then with E-cadherin-Fc (Sino Biological, Beijing, China). To obtain the first coating with protein A, 10 μL of the bead solution was added to a 1 mL PBS solution containing 5-10 mg EDC and 5-10 mg NHS, and 0.2 mg/mL protein A. The solution was allowed to incubate for 30 min at room temperature. The solution was then mixed in a 1 mL microcentrifuge tube on a shaker plate on a slow tilt to prevent sedimentation of the beads. The beads were then collected by using a pair of strong magnets (K&J Magnetics, Pipersville, PA) to pull the beads to the bottom of the tube and the supernatant solution was aspirated. Protein A functionalized beads were then resuspended in 400 μL of PBS with calcium, washed twice more, and then resuspended in 360 uL of PBS with calcium. Then, 40 uL of E-cadherin-Fc at 0.2 mg/mL was added. The solution was vortexed and incubated overnight at 4 °C. The beads were then washed with PBS with calcium three times and resuspended in 1 mL of PBS with calcium. The E-cadherin functionalized bead solution was stored at 4°C until further use.

### Magnetic Pulling Cytometer Setup

For force application we used a magnetic pulling cytometer fabricated with 2 strong neodymium magnets (91.25” diameter x 0.0625” thick from K&J Magnetics, Pipersville, PA, with a surface field of 662 Gauss) and an electrochemically sharpened and magnetically permeable 416 stainless steel probe positioned perpendicular to the magnets face, as described in our previous publication [64]. The magnets and probe were affixed by a 3D printed apparatus and attached to a 3-axis micromanipulator (ThorLabs) with a single motorized x-axis to move the probe towards or away from the bead.

### Probe Force Calibration and Bead Tracking

A silicone fluid (Sigma-Aldrich, ST. Louis, MO) with a viscosity of 1 Pa·s was added to the lid of a 60 mm cell culture dish and 10 μL of bead solution was added close to the base of the lid, manually stirred into the solution, and then placed in a vacuum chamber to degas for 30 min. The chamber, filled with the bead loaded silicone solution, was then setup onto the microscope stage and the magnetic pulling cytometer probe tip was immersed in this viscous medium with suspended magnetic beads. Magnetic beads were located and approached at a relative imaging plane similar to the setup when forces were applied to the cells, approximately 15 microns above the bead to prevent the probe tip from touching the coverslip. Images were captured every 0.33 s until the moving magnetic beads reached the probe. The beads were then tracked to calculate the instantaneous bead velocity at various distances from the probe which can be used to calculate the applied force using Stokes Law, F = 6πrηv [64, 65] where v is the instantaneous bead velocity, r is the magnetic bead radius and η is the medium viscosity (Fig. 1A). Bead motion was tracked using a custom MATLAB algorithm which located the bead using a reference image and normalized cross correlation of the images. For subpixel localization, images were scaled up 20-fold using the *imresize* function in MATLAB, which utilizes a bicubic interpolation.

**Figure 1.**
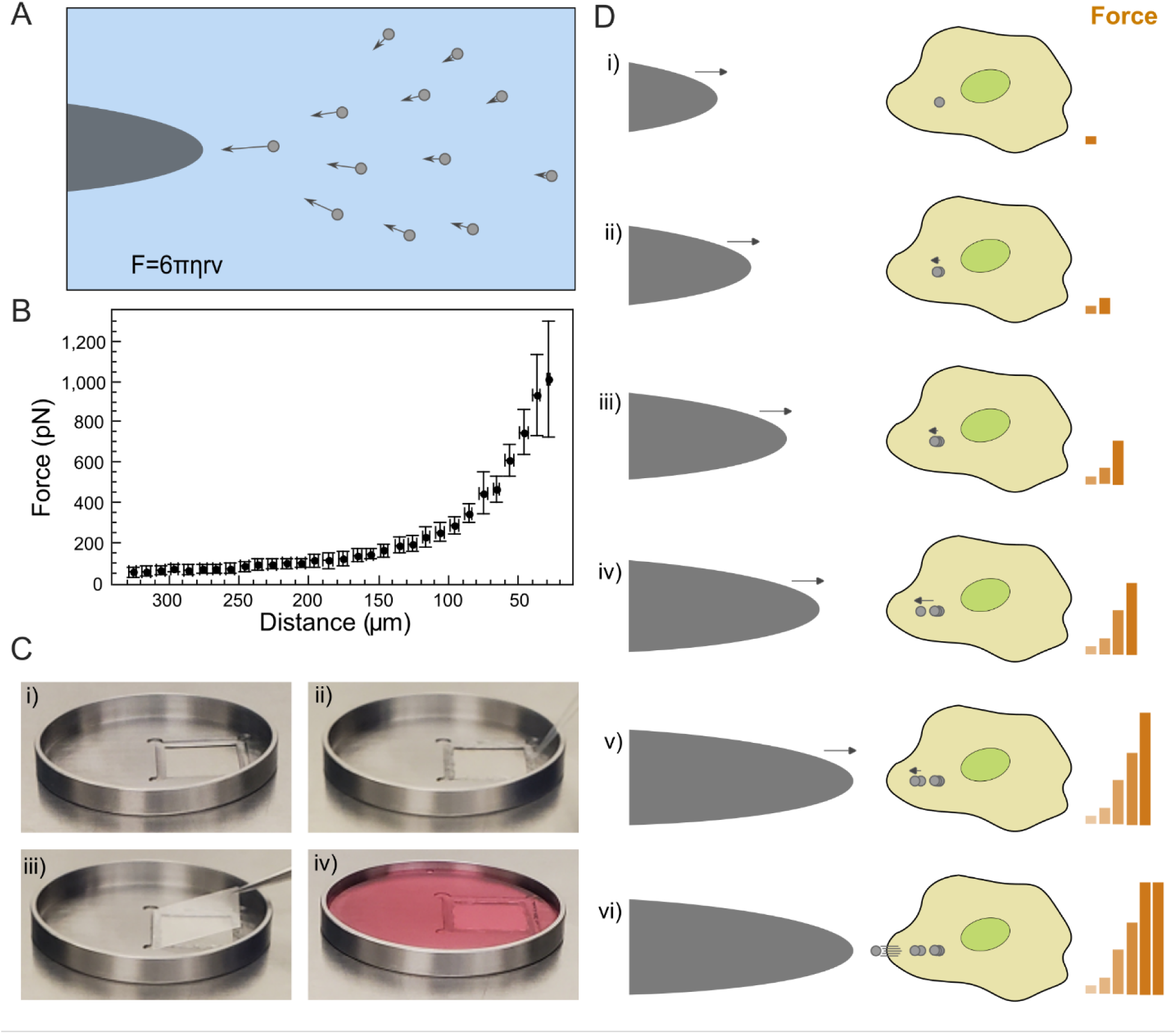
(A) Schematic depiction of calibration via the motion of microbeads through a viscous medium towards the magnetic probe. (B) Force versus distance calibration plot for the magnetic pulling cytometer (MPC). Error bars indicate the standard deviation around the mean of data points within 10-micron intervals. (C) Sample chamber used for experiments with beads on cells: (i) Stainless steel chamber with square shaped region to hold coverslip. (ii) Silicone grease was applied on the periphery of the square-shaped region. (iii) Glass coverslip (with plated cells and beads attached to the cells) was then affixed. (iv) Buffered cell culture media was then added. MPC probe was then made to approach from the left to apply forces on a chosen bead on a cell. (D) Schematic depiction of the exertion of sub-nanonewton forces on micron-scale E-cadherin adhesions between an E-cadherin coated 2.8 μm superparamagnetic bead and an epithelial cell using the magnetic pulling cytometer (MPC). The consistent incremental steps of the probe towards the bead implies a non-linear increase in loading rate due to the shape of the calibration curve in (A). The force applied at each step is obtained from the bead to probe tip distance and the non-linear increase in force is schematically depicted to the right of the cell. Tracking of the bead motion yields bead displacement and velocity as a function of time while the force applied at rupture gives the adhesion strength. Note that bead displacements are non-linear in time. At steps i-iii the bead displaces linearly, with a sudden rapid displacement between steps iii-iv, followed by bead deceleration between steps iv-v, and the bead detaches during the intermittent static hold of duration 1 second, between steps v-vi.

### Accelerated Force Ramp Application

Forces in the range of 100-800 pN were applied when the distance between the bead and probe tip was in the range of 315-40 μm (Fig. 1B). The force increases at an increasing rate with respect to the distance between the bead and the probe. The probe was set to approach at 5 μm intervals with a 1 second pause at each step. By approaching the bead at a constant rate, the magnitude of increase in force between steps, thus the loading rate itself increases such that an accelerated force ramp is applied.

### Force Application on Cells

A 316 stainless steel chamber was machined to replicate a 60 mm cell culture dish with a lower side wall height to increase the range of motion which could be achieved by the probe within the dish while avoiding contact with side wall. The chamber contained an 18 mm x 18 mm square opening with a lip (Fig. 1C) to affix the cell-plated coverslip with vacuum grease (Dow Corning, Midland, MI). Cell culture media with 10mM HEPES buffer was then added for imaging and force application. At 30 minutes prior to the experiment, 10 μL of the prepared E-Cadherin coated bead solution was pipetted onto the coverslip with the cells. The chamber and the magnetic pulling cytometer were then set up on the microscope stage and a cell-bound magnetic bead was located. The magnetic probe tip was positioned at a distance of 315 μm away from the magnetic bead, along a horizontal direction defined as the x-axis, and phase images of the cell and bead were acquired at 0.5s intervals until the bead-to-cell E-cadherin adhesion ruptured, which was observed by either the bead accelerating without ensuing deceleration or being immediately removed from the cell.

### Statistics

Pair-wise comparison of group means was performed with a t-test. One-way ANOVA was used to compare cell-cell contact phases amongst each other for WT as well as α-catenin KO cases. The symbols *, ** and *** were used to indicate p values <0.05, <0.01 and <0.001 respectively and n.s stands for not significant (p>0.05).

## Results and Discussion

To study the force response of E-cadherin adhesions, we used a custom-built magnetic pulling cytometer (MPC) set-up described previously [64]. We used the MPC to apply sub-nN scale forces on an E-cadherin-coated 2.8 μm superparamagnetic bead that was adherent upon an MDCK epithelial cell. We first calibrated the MPC by assessing the force exerted by the MPC probe tip on the superparamagnetic beads embedded in a silicone solution of known viscosity (Fig. 1A), as a function of distance as shown by the resulting force-distance calibration plot (Fig. 1B). As depicted in Figure 1B, for probe tip to bead distances greater than 40 μm, the force exerted stays below 1 nN. To enable use of the 40x oil immersion objective and expand the functional range of the probe tip within the sample, we designed a simple chamber machined of 316 Stainless Steel with a 60mm diameter (Fig. 1C) for use in the standard microscope housing. The overall measurement approach is schematically depicted in Figure 1D – as the MPC probe tip approaches at a constant rate, the bead attached to the cell via an E-cadherin adhesion experiences an applied force that increases at an accelerated rate and bead displacement is observed until adhesion failure. This protocol results not only in a non-linear increase in force, but also a non-linear increase in loading rate, the time derivative of the applied loading force. The applied force was determined from the probe-tip to bead distance and the calibration curve. The bead displacement was determined by tracking the E-cadherin bound bead as it moved along the cell resulting in the recording of force-dependent mobility of the E-cadherin adhesion. Finally, the adhesion strength was determined as the applied force when the E-cadherin bound bead detached from the actin cortex.

The non-linear increase in force and loading rate with time enables us to both observe the response of the bead, and therefore E-cadherin adhesion, to force as well as reach high enough forces to reach the adhesion strength and therefore observe the bead coming off the cell. Figure 2 shows phase images of a bead bound to a cell via an E-cadherin adhesion which is subject to increasing forces as the MPC probe tip approaches. Using the phase images, we tracked the paths taken by the E-cadherin-coated beads over time (Fig. 3A). There are some minor deviations in bead paths from the direction of force application presumably due to thermal motion as well as local obstructions on the cell surface along the bead path. After the last point on each path, the beads detached from the cell surface or began to accelerate and/or move at an increased linear rate until they detached. Since E-cadherin connects to the actin cytoskeleton via the catenins [66], and due to the previously reported role of α-catenin in modulating E-cadherin adhesion strength at the single molecule level [54] and in suspended cell doublets [56], we generated and employed MDCK cells in which α-catenin had been knocked out. Phase images of E-cadherin-coated beads bound to MDCK α-catenin KO cells subject to increasing forces as the MPC probe approaches, are shown in Figure 2B. The tracked E-cadherin bead paths on MDCK α-catenin KO cells are plotted in Figure 3Aii.

**Figure 2.**
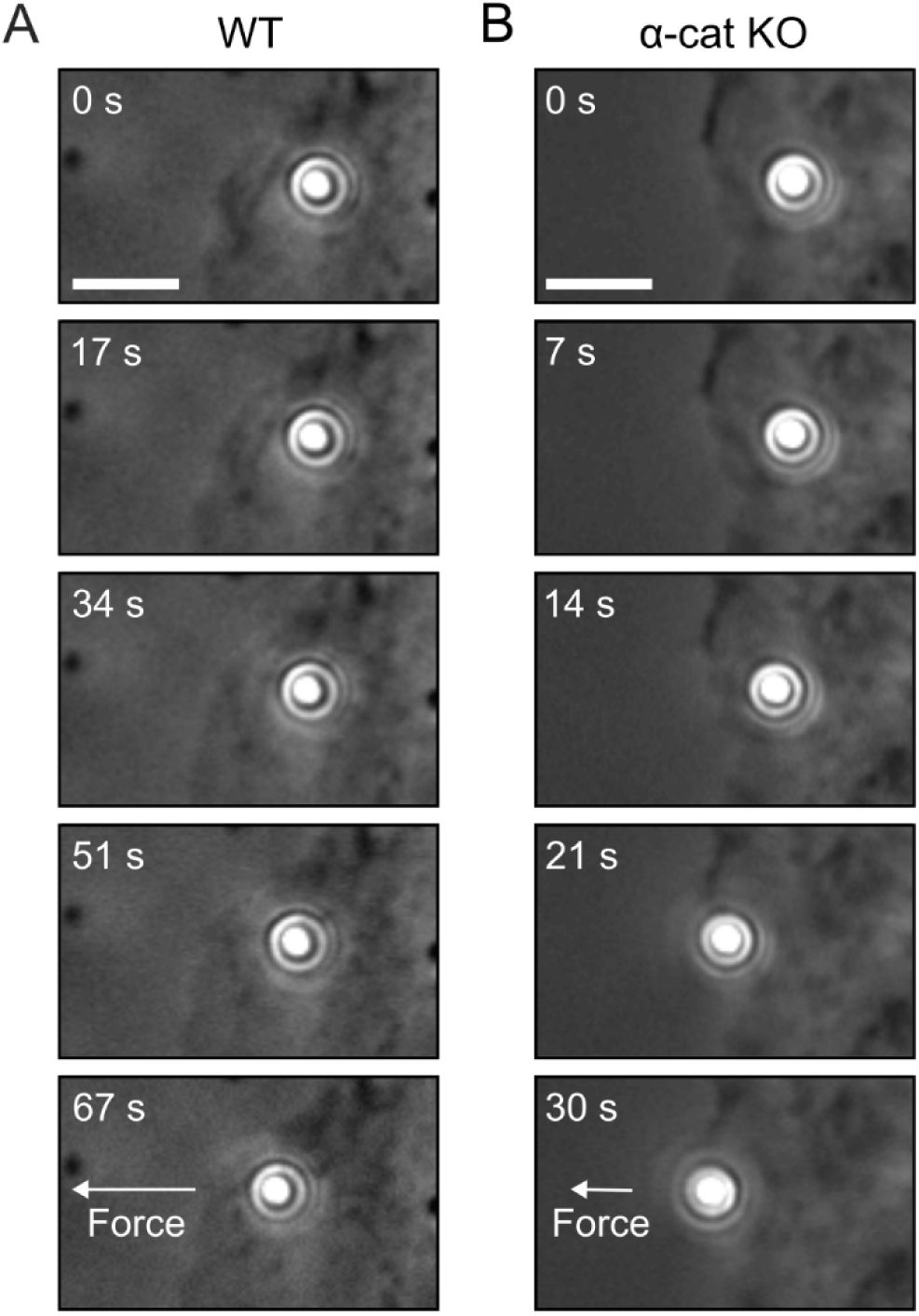
(A, B) Phase images of E-cadherin-coated beads attached to MDCK (A) or MDCK α-catenin KO (B) cells at five time points while increasing forces are applied via the MPC until adhesion rupture. Arrows in the bottommost panels indicate qualitative difference in force at time point before rupture. Scale bar is 5 μm.

**Figure 3.**
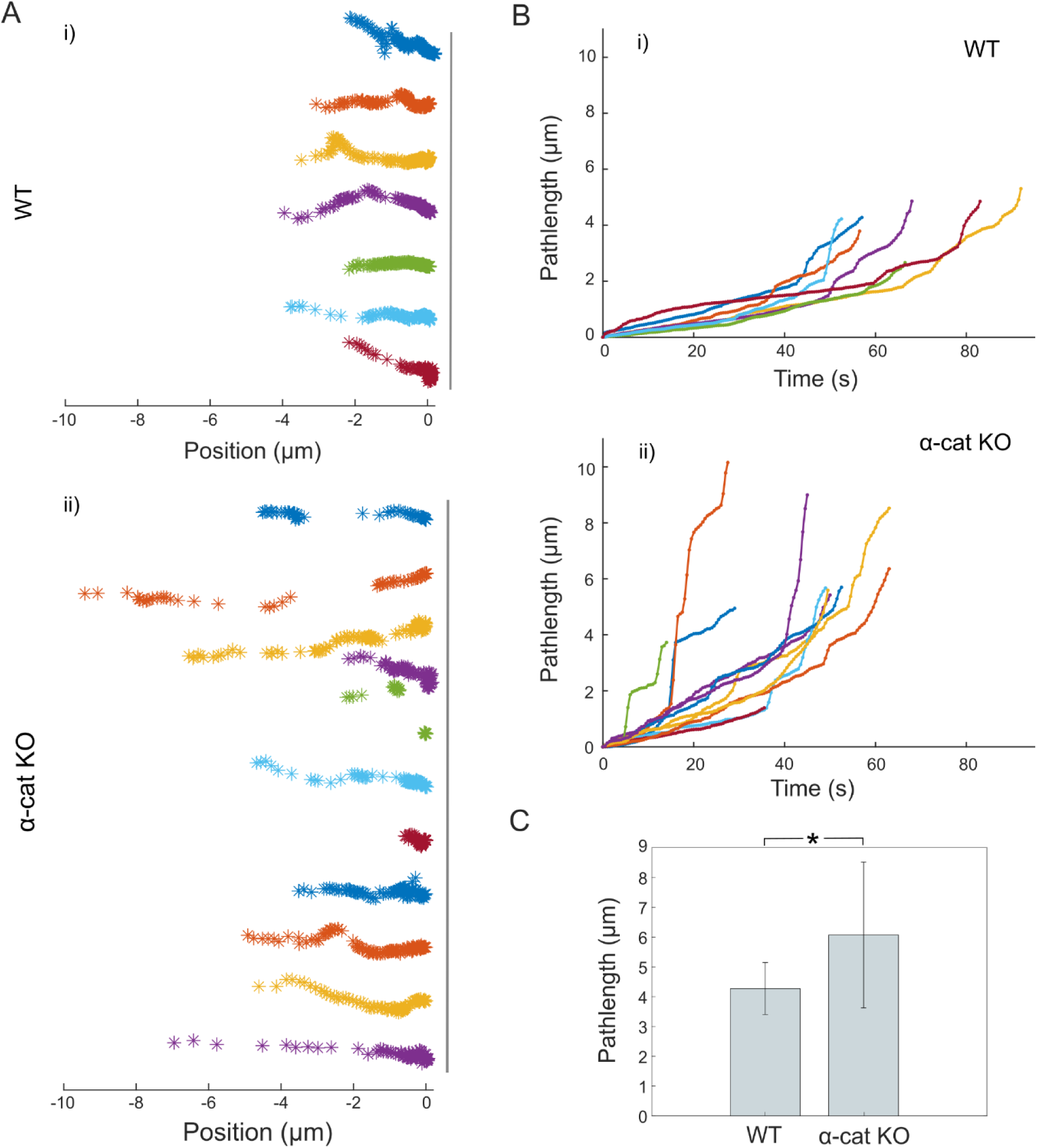
(A) Bead displacement tracks of E-cadherin-coated beads attached to MDCK (i) or MDCK α-catenin KO cells (ii) when subject to force. (B) Bead motion path length (when subject to force) as a function of time for MDCK (i) or MDCK α-catenin KO cells (ii). (C) Plot of the total bead path lengths of E-cadherin-coated beads attached to MDCK or MDCK α-catenin KO cells before adhesion rupture.

Comparison of E-cadherin bead paths on MDCK (Fig. 3Ai) and MDCK α-catenin KO cells (Fig. 3Aii) shows that the beads generally travelled farther on MDCK α-catenin KO cells than MDCK WT cells. Interestingly, comparison of the pathlength versus time for MDCK (Fig. 3Bi) and MDCK α-catenin KO cells (Fig. 3Bii) showed that the E-cadherin beads on MDCK α-catenin KO cells also travelled faster on average and had a larger mean pathlength (Fig. 3C). Careful examination of the plots of the pathlengths versus time (Fig. 3B) also showed that there are many instances of a relatively large displacement followed by the rate of increase in the pathlength, or velocity, reaching a local minimum. The deceleration during these sudden events indicates that the E-cadherin adhesion between the bead and the cell strengthens to resist force-induced motion, before continuing to slide at a relatively steady rate in response to increasing force until another such large displacement or rupture. Interestingly, these events appear to occur independently of any specific force level, or loading rate, suggesting this response may be the result of spontaneous ruptures within the local actin network, as opposed to force-dependent unbinding within the E-cadherin adhesion complex. The E-cadherin beads on the MDCK α-catenin KO cells also appeared to involve more sudden rapid displacements, which may be a result of a lower potential for intermittent adhesion strengthening, differing dynamics within the E-cadherin-to-actin complex, or the altered actin network structure as α-catenin dimers have been shown to compete with Arp2/3 for actin organization [44, 49, 67].

The adhesion strength i.e., the force required to rupture the adhesion and release the E-cadherin bead from its anchoring to the cell, was significantly lesser for beads on MDCK α-catenin KO cells than MDCK WT (Fig. 4A). Our measurement of adhesion strength at the micron-scale adhesion level is qualitatively consistent with previous results [54, 55] validating the importance of α-catenin for E-cadherin adhesion strength. We next wanted to assess if the force-dependent mobility of the E-cadherin adhesion between the bead and the cell, had any correlation to the adhesion strength. To first assess the resistance to motion of the bead-cell E-cadherin adhesion, we computed the ratio of the instantaneous force and the instantaneous bead velocity and averaged this ratio over the duration of bead motion to compute the effective drag coefficient. We found that the effective drag coefficient for the E-cadherin beads on MDCK α-catenin KO cells was significantly lesser than that on MDCK WT cells (Fig. 4B). A plot of the adhesion strength, from bead-cell E-cadherin adhesions, versus the effective drag coefficient of both MDCK and MDCK α-catenin KO cells taken together showed that these characteristics are positively correlated (Fig. 4C). This finding is consistent with the conceptualization that an adhesion with greater strength is more likely to couple strongly to the underlying actin cytoskeleton before rupture and thus this stronger coupling will result in greater resistance to adhesion motion in response to force.

**Figure 4.**
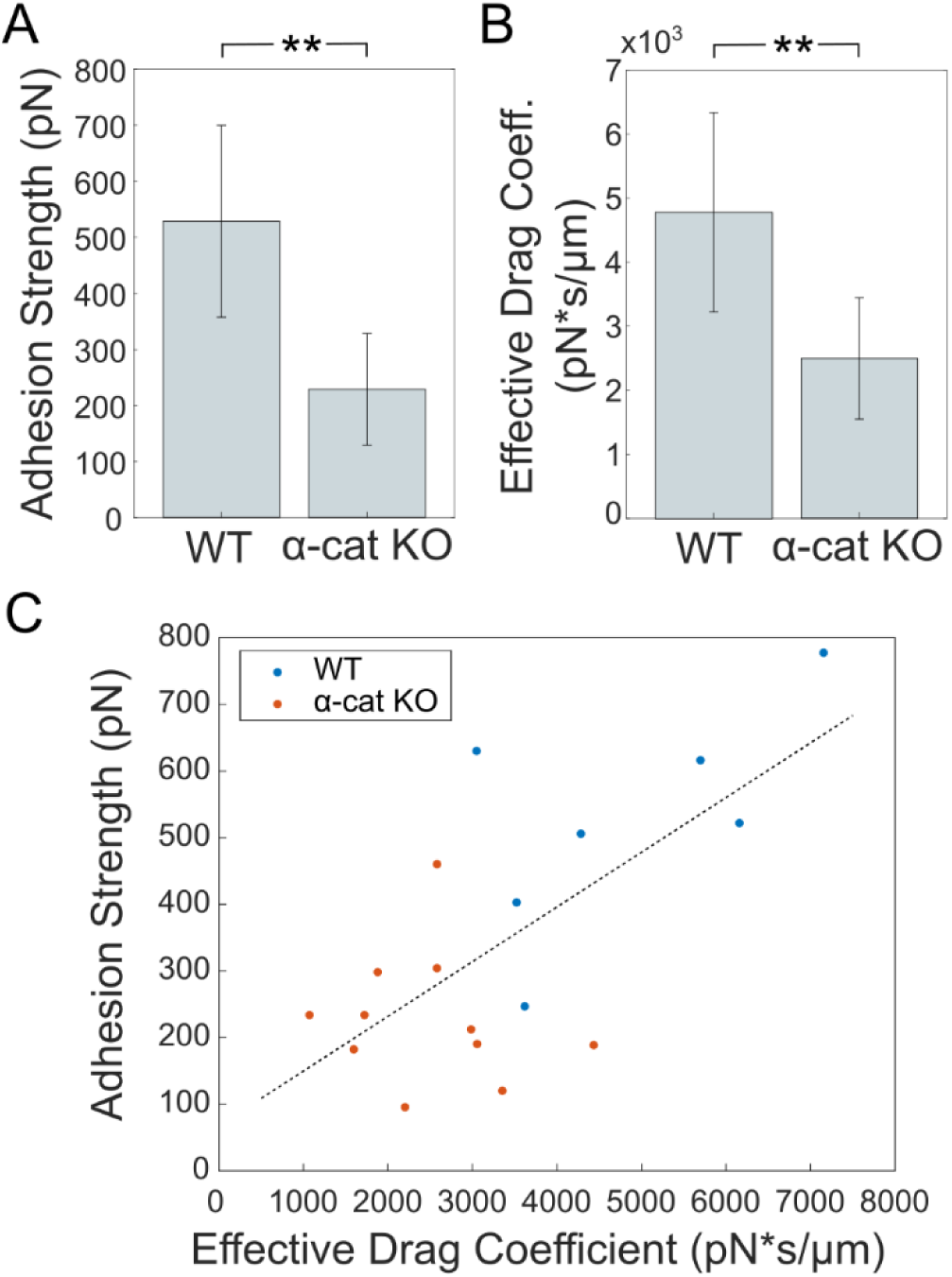
(A) Adhesion strength of E-cadherin adhesions between E-cadherin-coated beads and MDCK or MDCK α-catenin KO cells. (B) The average effective drag coefficient for E-cadherin adhesions between E-cadherin-coated beads and MDCK or MDCK α-catenin KO cells, computed as the average ratio of the instantaneous applied force and bead velocity till the adhesion ruptures. (C) Plot of the adhesion strength versus effective drag coefficient of the adhesion for all E-cadherin adhesions (MDCK and MDCK α-catenin KO cells), where the dotted line represents the linear trend.

Phase imaging of the cell-cell contacts of MDCK cells and MDCK α-catenin KO cells showed that the cell-cell contacts vary in morphology from smooth to fibrous when cells are pulling away, shown as phases (i-iv) in Figure 5A. We observed that the transition between these phases varied for WT cells and α-catenin KO cells. Specifically, WT cells showed a recovery of the cell-cell contact from the fibrous state (Video S1) while in α-catenin KO cells a fibrous state was maintained, or the contact separated (Video S2). By counting the frequency of each contact phase within cell islands, we observed that WT cells are most often smoother in morphology and rarely display a fibrous appearance (Fig. 5B). In contrast, cell-cell contacts for MDCK α-catenin KO cells showed a greater frequency of fibrous contacts and no significant difference in the frequency of each phase (Fig. 5B), demonstrating that α-catenin supports the maintenance of smooth cell-cell contacts in MDCK cells. Our observation that bead-cell E-cadherin adhesions lacking α-catenin exhibited more abrupt increases in bead motion under force, as well as the larger displacements, is consistent with the greater frequency of fibrous contacts. This suggests that these sudden internal rupture events enable the membrane bound E-cadherin adhesion to separate from the underlying actin cytoskeleton, whether at the adhesion complex-to-actin linkage or a deeper point within the network itself, and produce the observed fibrous contacts.

**Figure 5.**
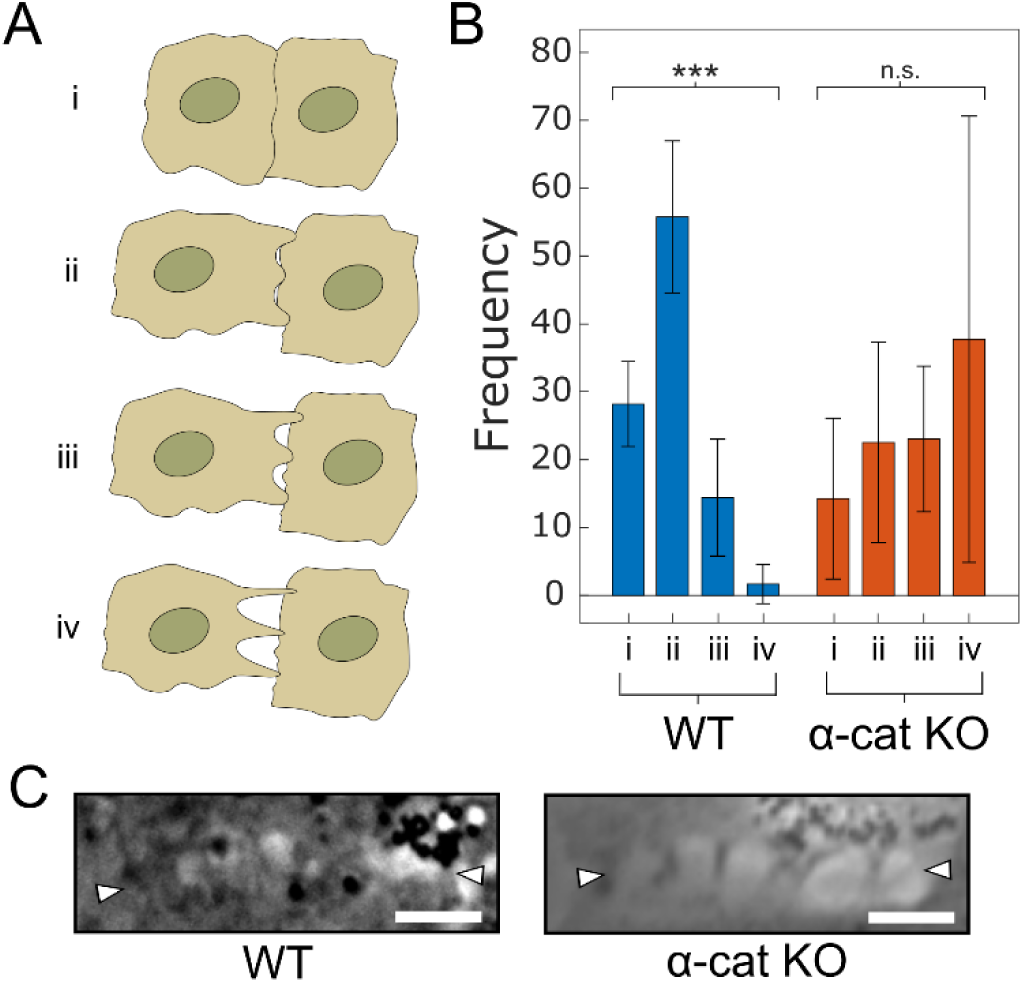
(A) Cartoon depiction of cell-cell contact transition from smooth to fibrous in different phases, (i) through (iv) (B) Plot of the frequency of cell-cell contacts in each phase (i-iv) between smooth and fibrous, for MDCK (WT) (from 7 cell islands and a total of 80 contacts) and MDCK α-catenin KO cells (from 7 cell islands and a total of 86 contacts). (C) Phase images of cell-cell contacts showing different fibrous contact morphologies for MDCK (WT) and MDCK α-catenin KO cells. White arrow heads indicate the cell-cell contact.

While fibrous cell-cell contacts in WT cells are still mostly contiguous, the fibrous contacts in the α-catenin KO cells display points of adhesion interspersed with gaps where the contact has intermittently ruptured (Fig. 5C). However, these gaps are not explained by the rapid displacement events of adhesions we observed in our bead-cell experiments. Namely because the adhesions can be preserved and reinforced after such events, as evidenced by the subsequent deceleration of adhesions. Notably, these rapid displacements often happened before bead-cell E-cadherin adhesion rupture in our experiments during which the applied magnetic force levels are maintained. However, within a true cell-cell adhesion, the applied force may be concomitantly diminished by deformation, contingent upon the force-dependent mobility of the adhesion. Thus, the lower effective drag coefficient observed in the α-catenin KO cells may permit the adhesions to more readily alleviate tension at the adhesion via an increased deformation rate acting as a negative feedback mechanism to preserve the adhesion in light of the lower adhesion strength. Without this negative feedback, the cell-cell contact would be more likely to rupture completely rather than producing a fibrous morphology. However, the lower adhesion strength may still cause strained adhesions to rupture more frequently and thus produce the observed gaps along the cell-cell contact. In sum, we propose that the spontaneous sliding events, in coordination with a decreased effective drag coefficient, enable the formation of sustained contact regions while the lower adhesion strength yields a lesser number of sustained contact regions creating the dynamic equilibrium state within the cell-cell contact. The unique nature of our force application protocol enabled us to observe adhesion motion in response to force in coordination with the eventual adhesion rupture while revealing sudden events of rapid displacement and reinforcement. The progressive increase in the force applied over time also implied a loading rate that was itself increasing with time. Therefore, the quantitative adhesion strengths and effective drag coefficients reported here are specific to our force application protocol which was used to compare the bead-cell E-cadherin adhesions on both MDCK and MDCK α-catenin KO cells.

## Conclusion

We have introduced a unique approach which we use to relate two key features of an adhesion – its force-dependent mobility before failure and its adhesion strength. We have demonstrated this approach here for individual E-cadherin adhesions between a micron-scale bead and an epithelial cell. We found that the resistance of the E-cadherin adhesion to force induced motion, captured by the effective drag coefficient, is positively correlated to its adhesion strength. This finding has important implications for understanding morphogenetic events at the cellular and adhesion scales, such as sustained cell-cell contact maintenance during multi-cellular rearrangements and migration. For instance, our results imply that weaker adhesions not only rupture at lower force levels but also displace at a faster rate in response to force. The correlation of these two properties implies that remodeling of cell-cell contacts can occur by adhesion sliding even if it does not eventually involve adhesion rupture. For example, by considering only adhesion strength, the default interpretation for contact remodeling is that more adhesions will be ruptured, but as a result of the correlation we observe, the weaker adhesions are also able to yield to a greater extent in response to lower force levels and thus reveals a negative feedback mechanism to preserve the adhesion longer, in opposition to the diminished adhesion strength, regulating cell-cell contact homeostasis. In addition to confirming the essential role of α-catenin in maintaining E-cadherin adhesion strength at the adhesion level, we also found that α-catenin specifically is also essential for sustaining a high effective drag coefficient displayed by the E-cadherin adhesion in response to force. It is tempting to speculate that this effect is due to α-catenin’s ability to couple to the actin cytoskeleton directly and through a plethora of linker molecules, however α-catenin’s concurrent role in actin organization warrants further study.

## Acknowledgments

V.M. acknowledges support from the National Institute of General Medical Sciences of the National Institutes of Health under award number 2R15GM116082. D.C. acknowledges support from the National Institute of General Medical Sciences of the National Institutes of Health under award number R35GM119617.

## References

1. Mandadapu, K.K., S. Govindjee, and M.R. Mofrad, On the cytoskeleton and soft glassy rheology. J Biomech, 2008. 41(7): p. 1467–78.

2. Ebata, H., et al., Activity-dependent glassy cell mechanics ?: Mechanical properties measured with active microrheology. Biophys J, 2023. 122(10): p. 1781–1793.

3. Kollmannsberger, P. and B. Fabry, Active soft glassy rheology of adherent cells. Soft Matter, 2009. 5(9).

4. Hang, J.-T. and G.-K. Xu, Stiffening and softening in the power-law rheological behaviors of cells. Journal of the Mechanics and Physics of Solids, 2022. 167: p. 104989.

5. Hang, J.T., et al., A hierarchical cellular structural model to unravel the universal power-law rheological behavior of living cells. Nat Commun, 2021. 12(1): p. 6067.

6. Mao, X. and Y. Shokef, Introduction to force transmission by nonlinear biomaterials. Soft Matter, 2021. 17(45): p. 10172–10176.

7. Hang, J.-T., G.-K. Xu, and H. Gao, Frequency-dependent transition in power-law rheological behavior of living cells. Science Advances, 2022. 8(18): p. eabn6093.

8. Chen, J., Nanobiomechanics of living cells: a review. Interface Focus, 2014. 4(2): p. 20130055.

9. Mao, Y., P. Nielsen, and J. Ali, Passive and Active Microrheology for Biomedical Systems. Frontiers in Bioengineering and Biotechnology, 2022. 10.

10. Mizuno, D., et al., Active and Passive Microrheology in Equilibrium and Nonequilibrium Systems. Macromolecules, 2008. 41(19): p. 7194–7202.

11. Robert, D., et al., In Vivo Determination of Fluctuating Forces during Endosome Trafficking Using a Combination of Active and Passive Microrheology. PLOS ONE, 2010. 5(4): p. e10046.

12. Chao, P.-Y., et al., Shear Wave Elasticity Measurements of Three-Dimensional Cancer Cell Cultures Using Laser Speckle Contrast Imaging. Scientific Reports, 2018. 8(1): p. 14470.

13. Caponi, S., et al., Non-contact elastography methods in mechanobiology: a point of view. European Biophysics Journal, 2022. 51(2): p. 99–104.

14. Larin, K., G. Scarcelli, and V. Yakovlev, Optical elastography and tissue biomechanics. J Biomed Opt, 2019. 24(11): p. 1–9.

15. Prevedel, R., et al., Brillouin microscopy: an emerging tool for mechanobiology. Nature Methods, 2019. 16(10): p. 969–977.

16. Kronenberg, N.M., et al., Long-term imaging of cellular forces with high precision by elastic resonator interference stress microscopy. Nature Cell Biology, 2017. 19(7): p. 864–872.

17. Hurst, S., B.E. Vos, and T. Betz, Intracellular softening and fluidification reveals a mechanical switch of cytoskeletal material contributions during division. bioRxiv, 2021: p. 2021.01.07.425761.

18. Weber, A., M.d. Vivanco, and J.L. Toca-Herrera, Application of self-organizing maps to AFM-based viscoelastic characterization of breast cancer cell mechanics. Scientific Reports, 2023. 13(1): p. 3087.

19. Weber, A., R. Benitez, and J.L. Toca-Herrera, Measuring biological materials mechanics with atomic force microscopy - Determination of viscoelastic cell properties from stress relaxation experiments. Microscopy Research and Technique, 2022. 85(10): p. 3284–3295.

20. Gallet, F., et al., Power spectrum of out-of-equilibrium forces in living cells: amplitude and frequency dependence. Soft Matter, 2009. 5(15): p. 2947–2953.

21. Wei, M.T., S.S. Jedlicka, and H.D. Ou-Yang, Intracellular nonequilibrium fluctuating stresses indicate how nonlinear cellular mechanical properties adapt to microenvironmental rigidity. Sci Rep, 2020. 10(1): p. 5902.

22. Puig-De-Morales, M., et al., Measurement of cell microrheology by magnetic twisting cytometry with frequency domain demodulation. Journal of Applied Physiology, 2001. 91(3): p. 1152–1159.

23. Evans, R.M.L., et al., Direct conversion of rheological compliance measurements into storage and loss moduli. Physical Review E, 2009. 80(1): p. 012501.

24. Aermes, C., et al., Environmentally controlled magnetic nano-tweezer for living cells and extracellular matrices. Scientific Reports, 2020. 10(1): p. 13453.

25. Mierke, C.T., et al., Focal adhesion kinase activity is required for actomyosin contractility-based invasion of cells into dense 3D matrices. Scientific Reports, 2017. 7(1): p. 42780.

26. Bidone, T.C., et al., Morphological Transformation and Force Generation of Active Cytoskeletal Networks. PLOS Computational Biology, 2017. 13(1): p. e1005277.

27. Yu, Q., et al., Balance between Force Generation and Relaxation Leads to Pulsed Contraction of Actomyosin Networks. Biophysical Journal, 2018. 115(10): p. 2003–2013.

28. Rossier, O.M., et al., Force generated by actomyosin contraction builds bridges between adhesive contacts. The EMBO Journal, 2010. 29(6): p. 1055–1068.

29. Chanet, S., et al., Actomyosin meshwork mechanosensing enables tissue shape to orient cell force. Nature Communications, 2017. 8(1): p. 15014.

30. Yang, Y.-A., et al., Local contractions regulate E-cadherin rigidity sensing. Science Advances, 2022. 8(4): p. eabk0387.

31. Feld, L., et al., Cellular contractile forces are nonmechanosensitive. Science Advances, 2020. 6(17): p. eaaz6997.

32. Perez-Vale, K.Z. and M. Peifer, Orchestrating morphogenesis: building the body plan by cell shape changes and movements. Development, 2020. 147(17).

33. Eftekharjoo, M., et al., Epithelial Cell-Like Elasticity Modulates Actin-Dependent E-Cadherin Adhesion Organization. ACS Biomater Sci Eng, 2022.

34. Hong, S., R.B. Troyanovsky, and S.M. Troyanovsky, Spontaneous assembly and active disassembly balance adherens junction homeostasis. Proceedings of the National Academy of Sciences, 2010. 107(8): p. 3528–33.

35. Borghi, N., et al., E-cadherin is under constitutive actomyosin-generated tension that is increased at cell–cell contacts upon externally applied stretch. Proceedings of the National Academy of Sciences, 2012. 109: p. 12568–12573.

36. Maruthamuthu, V., et al., Cell-ECM traction force modulates endogenous tension at cell-cell contacts. Proc Natl Acad Sci U S A, 2011. 108(12): p. 4708–13.

37. Maruthamuthu, V. and M.L. Gardel, Protrusive activity guides changes in cell-cell tension during epithelial cell scattering. Biophys J, 2014. 107(3): p. 555–63.

38. Dumbali, S.P., et al., Endogenous Sheet-Averaged Tension Within a Large Epithelial Cell Colony. J Biomech Eng, 2017. 139(10).

39. Buckley, C.D., et al., Cell adhesion. The minimal cadherin-catenin complex binds to actin filaments under force. Science, 2014. 346(6209): p. 1254211.

40. Wang, A., A.R. Dunn, and W.I. Weis, Mechanism of the cadherin-catenin F-actin catch bond interaction. Elife, 2022. 11.

41. Ishiyama, N., et al., Force-dependent allostery of the alpha-catenin actin-binding domain controls adherens junction dynamics and functions. Nat Commun, 2018. 9(1): p. 5121.

42. Hinck, L., et al., Dynamics of cadherin/catenin complex formation: novel protein interactions and pathways of complex assembly. J Cell Biol, 1994. 125(6): p. 1327–40.

43. Kim, T.J., et al., Dynamic visualization of alpha-catenin reveals rapid, reversible conformation switching between tension states. Curr Biol, 2015. 25(2): p. 218–24.

44. Desai, R., et al., Monomeric α-catenin links cadherin to the actin cytoskeleton. Nature Cell Biology, 2013. 15(3): p. 261–273.

45. Hazan, R.B., et al., Vinculin Is Associated with the E-cadherin Adhesion Complex. Journal of Biological Chemistry, 1997. 272(51): p. 32448–32453.

46. Itoh, M., et al., Involvement of ZO-1 in cadherin-based cell adhesion through its direct binding to alpha catenin and actin filaments. J Cell Biol, 1997. 138(1): p. 181–92.

47. Mandai, K., et al., Afadin: A Novel Actin Filament–binding Protein with One PDZ Domain Localized at Cadherin-based Cell-to-Cell Adherens Junction. Journal of Cell Biology, 1997. 139(2): p. 517–528.

48. Knudsen, K.A., et al., Interaction of alpha-actinin with the cadherin/catenin cell-cell adhesion complex via alpha-catenin. Journal of Cell Biology, 1995. 130(1): p. 67–77.

49. Kobielak, A. and E. Fuchs, Alpha-catenin: at the junction of intercellular adhesion and actin dynamics. Nat Rev Mol Cell Biol, 2004. 5(8): p. 614–25.

50. Abe, K. and M. Takeichi, EPLIN mediates linkage of the cadherin catenin complex to Factin and stabilizes the circumferential actin belt. Proc Natl Acad Sci U S A, 2008. 105(1): p. 13–9.

51. Yonemura, S., et al., alpha-Catenin as a tension transducer that induces adherens junction development. Nat Cell Biol, 2010. 12(6): p. 533–42.

52. Yao, M., et al., Force-dependent conformational switch of α-catenin controls vinculin binding. Nature Communications, 2014. 5(1): p. 4525.

53. Arbore, C., et al., α-catenin switches between a slip and an asymmetric catch bond with Factin to cooperatively regulate cell junction fluidity. Nature Communications, 2022. 13(1): p. 1146.

54. Saumendra Bajpai, J.C., Yunfeng Feng, Joana Figueiredo Sean X. Sun Gregory D. Longmore, and a.D.W. Gianpaolo Suriano, Alpha-Catenin mediates initial E-cadherin-dependent cell– cell recognition and subsequent bond strengthening. Proceedings of the National Academy of Sciences, 2008. 105: p. 18331–18336.

55. Thomas, W.A., et al., alpha-Catenin and vinculin cooperate to promote high E-cadherin-based adhesion strength. Journal of Biological Chemistry, 2013. 288(7): p. 4957–69.

56. Dufour, S., R.M. Mege, and J.P. Thiery, alpha-catenin, vinculin, and F-actin in strengthening E-cadherin cell-cell adhesions and mechanosensing. Cell Adh Migr, 2013. 7(4): p. 345–50.

57. Cavey, M., et al., A two-tiered mechanism for stabilization and immobilization of Ecadherin. Nature, 2008. 453(7196): p. 751–6.

58. Xie, B., et al., Molecular mechanism for strengthening E-cadherin adhesion using a monoclonal antibody. Proc Natl Acad Sci U S A, 2022. 119(32): p. e2204473119.

59. Chu, Y.S., et al., Force measurements in E-cadherin-mediated cell doublets reveal rapid adhesion strengthened by actin cytoskeleton remodeling through Rac and Cdc42. J Cell Biol, 2004. 167(6): p. 1183–94.

60. Tabdili, H., et al., Cadherin-dependent mechanotransduction depends on ligand identity but not affinity. J Cell Sci, 2012. 125(Pt 18): p. 4362–71.

61. Muhamed, I., F. Chowdhury, and V. Maruthamuthu, Biophysical Tools to Study Cellular Mechanotransduction. Bioengineering, 2017. 4(1): p. 12.

62. le Duc, Q., et al., Vinculin potentiates E-cadherin mechanosensing and is recruited to actin-anchored sites within adherens junctions in a myosin II-dependent manner. Journal of Cell Biology, 2010. 189(7): p. 1107–15.

63. Ozawa, M., Nonmuscle myosin IIA is involved in recruitment of apical junction components through activation of alpha-catenin. Biol Open, 2018. 7(5).

64. Bush, J. and V. Maruthamuthu, In situ determination of exerted forces in magnetic pulling cytometry. AIP Adv, 2019. 9(3): p. 035221.

65. Andreas R. Bausch, W.M., Erich Sackmann, Measurement of Local Viscoelesticity and Forces in Living Cells by Magnetic Tweezers. Biophysical Journal, 1999. 76: p. 573–579.

66. Gumbiner, B.M., Regulation of cadherin-mediated adhesion in morphogenesis. Nature Reviews in Molecular and Cell Biology, 2005. 6(8): p. 622–634.

67. Drees, F., et al., Alpha-catenin is a molecular switch that binds E-cadherin-beta-catenin and regulates actin-filament assembly. Cell, 2005. 123(5): p. 903–15.

