## Supplementary Video Legend for "α-Catenin Dependent E-cadherin Adhesion Dynamics as Revealed by an Accelerated Force Ramp"

### Supplementary Material

**Supplementary Video S1.** Timelapse video of MDCK WT cells over a 30-minute duration. A central cell is observed to spontaneously contract while the lower right contact transitions into a fibrous state and then recovers to a smooth state by local relaxation of the neighboring cell. The neighboring contacts are able to sustain a relatively smooth contact state during this event and initiate strains within the neighboring cells themselves, as opposed to alleviating the stress by significant strains, and resulting fiber, within the contact itself. At the lower left the central cell sustains a singular long fiber across neighboring cells and notably sustains enough tension to alter the local geometry.

**Supplementary Video S2.** Timelapse video of MDCK α-catenin KO cells over a 30-minute duration. At the lower region a cell spontaneously contracts and separates from its neighboring cell contacts while maintaining long fiber contacts. Two cells/contacts above this one, there is a fibrous contact that is maintained and somewhat begins to return to a smoother state as the cell to the right of the contact relaxes. The contact above this one (upper right) shows a cell contact transiently separating as the generated fibrous contacts also rupture. Notably these fibers do not show significant deformation at the local contact point as the fibers thin out until rupture.
